# Boosting Toll-like receptor 4 signaling enhances the therapeutic outcome of antibiotic therapy in pneumococcal pneumonia

**DOI:** 10.1101/2020.02.18.955500

**Authors:** Fiordiligie Casilag, Sebastian Franck, Laura Matarazzo, Martin Figeac, Robin Michelet, Charlotte Kloft, Christophe Carnoy, Jean-Claude Sirard

**Affiliations:** Univ. Lille, CNRS, INSERM, CHU Lille, Institut Pasteur de Lille, U1019 – UMR9017 - CIIL - Center for Infection and Immunity of Lille, F-59000 Lille, France; Freie Universitaet Berlin, Institute of Pharmacy, Dept. of Clinical Pharmacy & Biochemistry, D-12169 Berlin, Germany; Univ. Lille, Plateforme de Génomique Fonctionnelle et Structurale, F-59000 Lille, France

## Abstract

The emergence and spread of antibiotic resistance emphasize the need for alternative treatment strategies against bacterial infections. Boosting the host innate immunity is not only readily deployable in most individuals but can also mobilize many different antibacterial defenses. This study tested the hypothesis whereby stimulation of the innate immune receptor Toll-like receptor 4 (TLR4) can be combined with antibiotics in the treatment of invasive pneumonia. In a mouse model of *Streptococcus pneumoniae* infection, a single oral administration of low-dose amoxicillin (AMX) or the systemic delivery of monophosphoryl lipid A (MPLA, a clinically-approved TLR4 activator) decreased the bacterial load in lung and spleen, although this was not sufficient for long-term survival. In contrast, a single treatment with a combination of MPLA and AMX induced significant bacterial clearance with little to no regrowth over time, and was associated with longer survival. Upregulation of genes related to granulocyte infiltration in lung tissue and elevation of blood levels of pro-inflammatory cytokines was immediate and transient in MPLA-treated mice; this indicates activation of the innate immune system in a context of infection. Combination treatment was associated with a well-preserved lung tissue architecture and more rapid recovery from inflammation - suggesting that immune activation by MPLA does not exacerbate pneumonia-induced damage. After AMX administration, plasma AMX concentrations rapidly reached the maximum and declined, whereas the downstream effects of MPLA extended beyond AMX elimination; these findings suggested a two-step effect. Our results demonstrated that leveraging host innate immunity increases the efficacy of antibiotic therapy in bacterial pneumonia.

## INTRODUCTION

The discovery and development of antibiotics in the 20^th^ century was a major turning point in medicine; it enabled the successful treatment and/or prevention of many infectious diseases in humans and in other animals. Decades later, these drugs are back in the headlines but for the wrong reasons: the alarming decline in their therapeutic effectiveness and the spread in antimicrobial resistance (AMR). The latter is a major threat to human health because it compromises our ability to treat bacterial infections and to carry out medical procedures that rely on prophylactic antibiotic use, such as chemotherapy, transplantation, and surgery (1). A single pathogen can express multiple resistance mechanisms, which in turn can often confer protection against several classes of antibiotics. In a clinical setting, this usually necessitates treatment with a “last resort” antibiotic or with combinations of antibiotics. In 2015, a report from the World Health Organization raised concerns about the lack of new antibiotics in development (2). This emphasizes the need to come up with innovative anti-infective approaches for treating resistant pathogens and preventing the further dissemination of AMR.

Advances in medicine and technology have provided deeper insights into immunology and thus increased the viability of host-directed therapeutic strategies. For example, the targeted stimulation of innate immunity using immunomodulatory drugs has gained much attention in recent years (3). This approach has three main advantages: (i) it makes use of universal, built-in machinery that is ready to activate in most individuals, (ii) it has both anti-infective and pro-recovery effects, and (iii) the complex innate immune system mobilizes many different effectors through a tightly coordinated string of events, and thus counters the potential development of AMR by invading pathogens. The host’s recognition of immediate danger is instrumental in mounting a successful defense against invading pathogens. *In vivo*, many cell types are equipped with pattern-recognition receptors; for example, Toll-like receptors (TLRs) are present on many cell types and are capable of binding to conserved macromolecules expressed by microorganisms (4). Thus, TLR engagement by an agonist triggers signaling cascades that lead to the transcriptional activation of immune genes and the regulation of antibacterial mechanisms for eliminating the threat. Signaling by TLRs triggers the production of various chemokines and antimicrobial compounds, activates complement, and stimulates leukocyte differentiation and mobilization. In view of the TLR-dependent response mechanisms’ early involvement in host defense, their robust activation and their highly inducible nature, researchers have sought to design novel or improved current therapies against viral and bacterial infections (3, 5). Monophosphoryl lipid A (MPLA) is a derivative of the immunostimulatory lipid A component of the outer-membrane-expressed lipopolysaccharide (LPS) from the bacterium *Salmonella minnesota* R595 (6). Lipopolysaccharide itself is highly toxic, due to its ability to strongly activate TLR4 downstream signaling at low doses via both of the receptor’s adaptor proteins, namely myeloid differentiation primary response 88 (MyD88) and Toll/interleukin-1 receptor domain-c activation protein inducing interferon beta (TRIF). In contrast, activation of TLR4 by MPLA is biased towards TRIF-dependent TLR4 signaling, making MPLA safe for use in humans (7). Monophosphoryl lipid A induces a significant but attenuated innate immune response (8-10); this property is related to the differences in its molecular structure vs. LPS (11, 12). The combination of immunostimulatory activity and low toxicity make MPLA an attractive candidate for therapeutic use in humans. Years of research have paved the way to MPLA becoming the first TLR agonist to be licensed as an adjuvant in certain vaccine formulations (12-14). Despite the growing body of literature data (indicating continued interest in finding further applications for MPLA), most studies have focused on its use as a prophylactic treatment. Indeed, MPLA has been shown to confer protection against infections by (i) *Pseudomonas aeruginosa* in burn-wounds, (ii) *Staphylococcus aureus* under post-hemorrhagic conditions, and (iii) nontypeable *Haemophilus influenzae* in the nasopharynx (15-18). In view of these observations, we hypothesized that MPLA may be a viable treatment against an ongoing bacterial infection, and so looked into both its applicability and efficacy as an anti-infective therapy. To the best of our knowledge, the only other previous study of this approach was performed in the context of fungal infection (19).

In order to tackle the challenges of developing alternative therapeutic strategies against bacterial infections and combating the spread of AMR, we designed and performed the present proof-of-concept study. The prime objective was to establish whether host immune responses can be leveraged to achieve a successful treatment outcome. Using a previously established murine model of invasive pneumococcal disease (20, 21), we determined whether deliberately TLR4-activated innate immune responses can constitute an adjunct to standard antibiotic therapy, improve the latter’s efficacy, and/or promote quicker tissue recovery after an infection. To this purpose, the study investigates the MPLA effect on amoxicillin (AMX), a beta-lactam antibiotic used as first-line treatment against *S. pneumoniae*.

## RESULTS

### The combination of MPLA and amoxicillin increases *S. pneumoniae* clearance and extends survival

Intraperitoneal administration of MPLA to naïve animals (0.5 to 50 µg per mouse) increased the mRNA and protein levels of inflammatory mediators - indicating MPLA’s ability to induce innate immune responses three hours post-administration (**Figure S1**). The magnitude of the systemic immune responses (i.e. in the liver and blood) was strongly dependent on the dose of MPLA. Innate immune responses were also observed in the lungs after the systemic injection of MPLA, albeit only at the highest dose (50 µg). This result suggests that the systemic administration of MPLA promotes both systemic and pulmonary immune responses - a feature that could potentially be exploited in the host-directed therapy of respiratory infectious diseases.

We next looked at whether MPLA’s immunomodulatory effects impacted the bacterial load during pneumococcal infection in mice when the TLR4 activator was administered together with AMX, a first-line treatment against *S. pneumoniae*. Twelve hours after intranasal inoculation with *S. pneumoniae*, Swiss (CD-1) mice received either a sub-curative dose of AMX (10 µg per animal; 0.4 mg/kg, administered by oral gavage), MPLA (50 µg per animal; 2.0 mg/kg, administered by intraperitoneal injection) or a combination of the two treatments (AMX+MPLA). Bacterial counts in the lungs and spleen were determined at different time points as surrogate markers of pneumonia and bacterial dissemination, respectively (**Figure 1**). At 24 hours post-infection, the bacterial loads in the lungs and spleens were lower in all treated animals than in mock-treated animals; however, the differences between the three treatments were not statistically significant (**Figure 1B-D**). In contrast, we observed significant intergroup differences in the bacterial loads at 48 hours post-infection (**Figure 1B-C and 1E**). Bacterial clearance was greatest in the AMX+MPLA group, with a median CFU value of 5.1 × 10^2^ in the lungs, and nearly undetectable bacterial levels in the spleen; this can be compared with values of 3.6 × 10^7^ and 1.9 × 10^6^ CFU recorded in the lungs and spleen of mock-treated infected animals, respectively (corresponding to 7.1 × 10^4^- and 1.1 × 10^5^-fold differences in the bacterial load, respectively). The lung bacterial load was 1.3 × 10^5^ CFU for AMX-treated mice and 9.4 × 10^3^ CFU for MPLA-treated mice, i.e. respectively 255 and 18 times higher than in the AMX+MPLA group. Importantly, AMX and MPLA monotherapies were unable to prevent bacteremia; the respective median splenic bacterial loads were 4.6 × 10^3^ and 1.8 × 10^2^ CFU.

**Figure 1.**
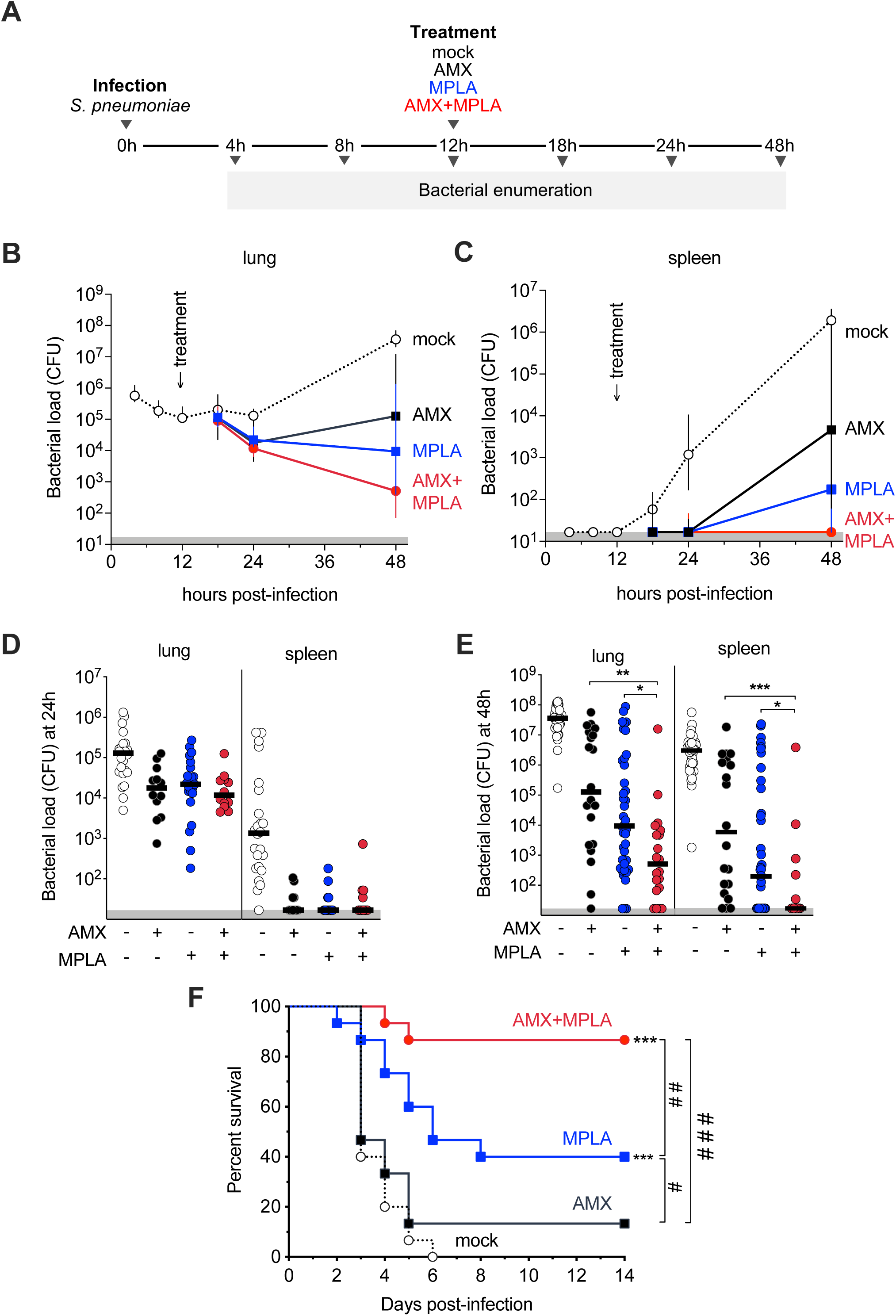
Combination treatment with AMX and MPLA is effective against *S. pneumoniae in vivo*. (A) CD-1 mice were infected intranasally with 1×10^6^ *S. pneumoniae* and then given either 10 µg of AMX intragastrically, 50 µg MPLA intraperitoneally, a combination of the two treatments (AMX+MPLA), or water and saline mock treatments 12 hours post-infection. Lungs and spleens were collected at different time points for quantification of the bacterial load using standard plate counting methods. (B-C) Bacterial growth over time in infected mice, showing the total bacterial load in the indicated tissues (as CFUs). Symbols represent the median value (n≥6/group) and error bars represent the interquartile range; the gray shaded area along the x-axis indicates the assay’s limit of detection. (D-E) Lung and spleen bacterial counts from individual mice (n≥12/group) 24 h (D) and 48 h (E) post-infection. The solid lines indicate the median value for each group, and the gray shaded area along the x-axis indicates the assay’s limit of detection. A one-way ANOVA (the Kruskal-Wallis test with Dunn’s post-test for multiple comparisons) was applied. *=p<0.05, **=p<0.01, ***=p<0.001 vs. the indicated comparator groups. Data from the mock control group (shown in white) were excluded from statistical analyses of treatment groups. (F) Survival curves (n=15 mice per group). Gehan-Breslow-Wilcoxon test. *** = *p*<0.001 vs. infected mock treated control, # = p<0.05, # # = *p*<0.01, # # # = *p*<0.001 vs. the indicated comparator groups.

Interestingly, the bacterial load in the spleen was a strong predictor of survival (**Figure 1F**). All mock animals succumbed to infection within 3 to 6 days, whereas AMX and MPLA monotherapies were associated with survival rates of 13.3% and 40%, respectively. The AMX+MPLA treatment outperformed the two monotherapies, with a survival rate of 86.7% - more than twice the value for MPLA, and over six times the value for AMX. It is noteworthy that the difference in the survival rate between AMX+MPLA treatment and high-dose AMX monotherapy (30 µg per animal; 1.2 mg/kg) was not significant (**Figure S2A**). This suggests that co-administration of MPLA with low-dose AMX can boost the antibiotic’s efficacy to levels comparable with standalone, higher-dose treatment. A similar potentiating effect was observed in the congenic BALB/c mice (**Figure S2B**). Overall, the present results demonstrate that AMX+MPLA combination treatment improves the therapeutic outcome of low-dose AMX and is efficacious against *S. pneumoniae in vivo* by minimizing bacterial lung colonization and dissemination, and promoting long-term survival.

### Combination treatment with AMX+MPLA mitigates pneumonia-induced lung damage

We next looked at whether or not MPLA-mediated pro-inflammatory signaling exacerbated inflammation due to *S. pneumoniae* infection. To this end, the lung tissue architecture in animals having been treated 12 hours post-infection with AMX, MPLA or AMX+MPLA was analyzed 48 hours post-infection. As a positive control, a group of animals was treated with a single, curative, high dose of AMX (350 µg per animal; 14 mg/kg). The histopathological assessment revealed that treatment with MPLA in the presence or absence of AMX did not exacerbate lung inflammation (**Figure 2**). Notably, all the AMX+MPLA-treated mice did not show any signs of the perivascular inflammatory cell infiltration observed in the other groups (including animals having receiving the curative dose of AMX). The total histopathological scores also showed that AMX+MPLA treatment had the greatest impact on preservation of the lung tissue architecture, with the lowest score of 5; the corresponding values were 5.75, 6.25, 8.75 and 12 in the high-dose AMX, MPLA, low-dose AMX and mock groups, respectively (**Figure 2F)**. These findings suggest that MPLA treatment not only mitigate the effects of infection-induced tissue damage but also (when combined with AMX) promotes tissue recovery and improves the antibiotic’s efficacy without exacerbating inflammation.

**Figure 2.**
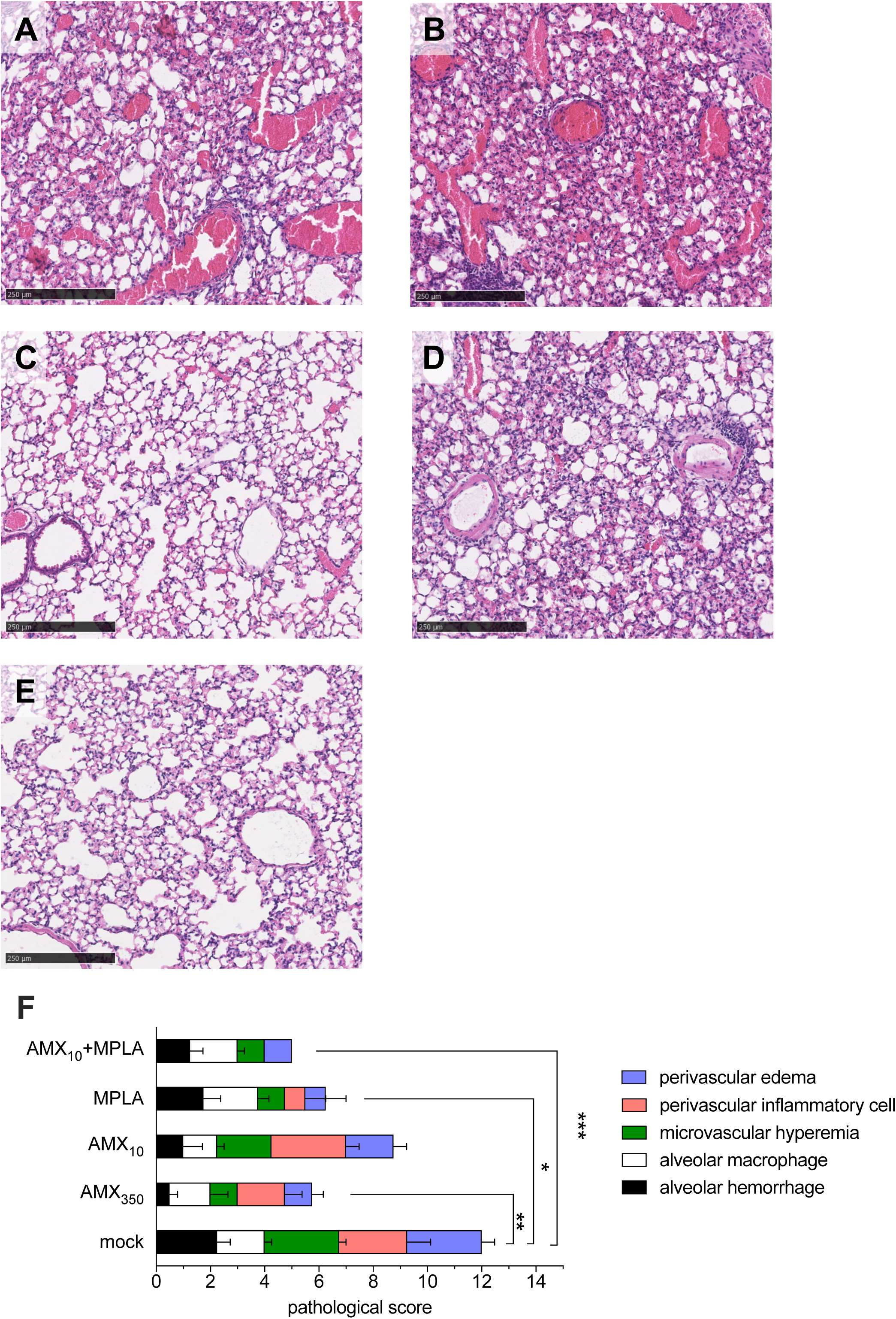
Combination treatment with AMX and MPLA mitigates infection-induced tissue damage. CD-1 mice (n=4/group) were infected intranasally with 1×10^6^ *S. pneumoniae*, and then given treatments 12 hours later as indicated: (A) No treatment, i.e. mock, (B) intragastric treatment with 10 µg of amoxicillin (AMX_10_) or (C) 350 µg amoxicillin (AMX_350_), (D) intraperitoneal treatment with 50 µg of MPLA (MPLA), or (E) a combination of intragastric treatment with 10 µg AMX and intraperitoneal treatment with 50 µg of MPLA (AMX+MPLA). Hematoxylin- and eosin-stained tissue sections showing the lung architecture 48 hours post-infection (A-E). The images are representative of four biological replicates per group. Scale bar = 150 µm. (F) Histopathological scores were assessed on a 0-5 scale: 0=absence, 1=minimal, 2=slight, 3=moderate, 4=marked, and 5=severe. The bars represent the mean ± SEM. A one-way ANOVA (the Kruskal-Wallis test with Dunn’s post-test for multiple comparisons) was applied. *=p<0.05 and **=p<0.01.

### Systemic MPLA treatment boosts the airway’s innate immune responses during pneumonia

It has already been shown that MPLA confers protection against bacterial challenge when administrated prophylactically, i.e. prior to the infectious challenge (15, 17, 18). Our study showed that MPLA also has an immunomodulatory effect after the infection has been established. To further characterize the local immune responses that were elicited and could participate to bacterial clearance, we used microarrays to analyze the transcriptome of lung tissue. We then investigated post-treatment changes in gene expression by initially comparing the AMX- and AMX+MPLA-treated groups 2, 4, and 8 h after treatment (i.e. 14, 16 and 20 h post-infection) (**Figure 3**). The overall response to the AMX+MPLA treatment indicated an enrichment in the granulocyte adhesion and diapedesis pathway and the leukocyte mobilization pathways (**Figure 3B**). Moreover, the difference between the AMX- and AMX+MPLA-treated groups in the number of transcripts that were expressed >2- or <0.5-fold was highest at 2 h post-treatment (n=188 transcripts) and decreased over time, with 106 transcripts at 4 h and 13 transcripts at 8 h (**Figure 3C**). There were 173 upregulated transcripts at 2 h, 75 at 4 h, and 12 at 8 h (**Figure 3C**). The pattern and time course of expression in lungs suggested that the MPLA-induced transcriptional effects at the infection site were immediate and transient. Some of the lung transcripts strongly expressed within a few hours of treatment were associated with neutrophil function (e.g. *Ngp, Itgb2l*, and *Mmp8*) or encoded proteins with known antibacterial properties (e.g. CAMP or S100A8) (**Figure 3D**).

**Figure 3.**
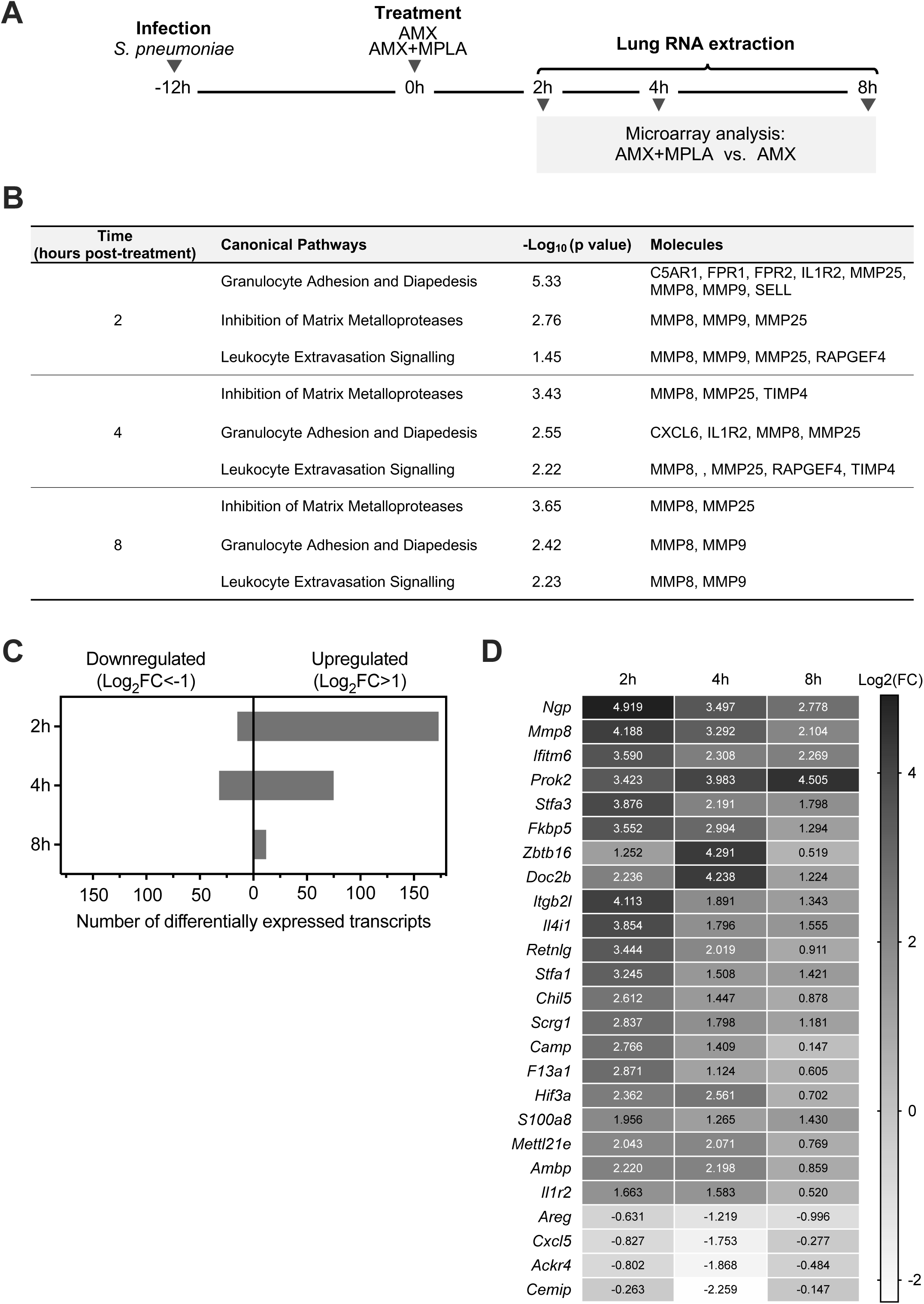
Characterization of local immune response signatures of MPLA treatment. (A) BALB/c mice (n=3/group) were infected with 1×10^6^ *S. pneumoniae* and then treated with either 10 µg of intragastrically administered AMX or a combination of AMX and 50 µg of intraperitoneally administered MPLA (AMX+MPLA). Total RNA was extracted from lungs collected at different time points, and mRNA transcripts were compared in a microarray analysis. (B) Enrichment of canonical pathways, according to an Ingenuity Pathway Analysis of the microarray datasets. (C) The number of transcripts with significantly greater expression (upregulation, Log_2_(fold change [FC] >1) or significantly lower expression (downregulation, Log_2_FC < -1) in (AMX+MPLA)-treated vs. AMX-treated animals. (D) The 25 genes with the highest or lowest differential expression levels in the microarray analysis (AMX+MPLA vs. AMX);

In a series of follow-up experiments *in vivo*, we confirmed the above results by using RT-qPCR assays for selected genes. We extended the transcriptional study by including groups of infected and untreated animals, infected and MPLA-treated animals, and uninfected and untreated (naïve) animals (**Figure 4**). In line with the microarray data, MPLA treatment was found to accelerate the onset of potentially antimicrobial-related *Ngp, Itgb2l, Mmp8, Camp, S100a9, Fkbp5, Ifitm6, Il4i1, Prok2*, and *Zbtb16* transcript expression in infected animals in both the MPLA-only and AMX+MPLA groups (**Figure 4B**). The effect of MPLA on transcript expression was therefore independent of AMX treatment. The lung expression of pro-inflammatory genes coding for cytokines and chemokines (such as *Ccl2, Ccl20, Il1b*, or *Il12b*) was increased by infection but was not further impacted by MPLA or AMX+MPLA treatment - indicating that the expression of these genes was primarily regulated in an infection-dependent manner (**Figure 4B**). Given that we compared AMX+MPLA-treated with AMX-treated groups in our initial screening, this might explain why we failed to detect the differential expression of *Ccl2, Ccl20, Il1b*, or *Il12b* in our microarray experiments. Lastly, by comparing infected and untreated animals, infected and MPLA-treated animals, and uninfected and MPLA-treated animals (**Figure S3**), we found that the transcription of some genes was somehow dependent on both infection- and MPLA-induced signaling; this suggests the presence of a priming effect in which prior *S. pneumoniae* infection results in a more robust response following the administration of MPLA. We also found that the AMX+MPLA combination did not influence the neutrophil count in the lungs and spleen 12 h post-treatment (**Figure S4**). A similar pattern was observed for alveolar macrophages. In contrast, the lung monocyte count was higher in the AMX+MPLA group than in the mock treatment group (**Figure S4A**). Overall, these results demonstrated that while a bacterial pneumonia insult can prompt airway innate immune responses, the latter are enhanced by post-infection treatment with MPLA; this probably contributes to greater bacterial clearance in the lungs.

**Figure 4.**
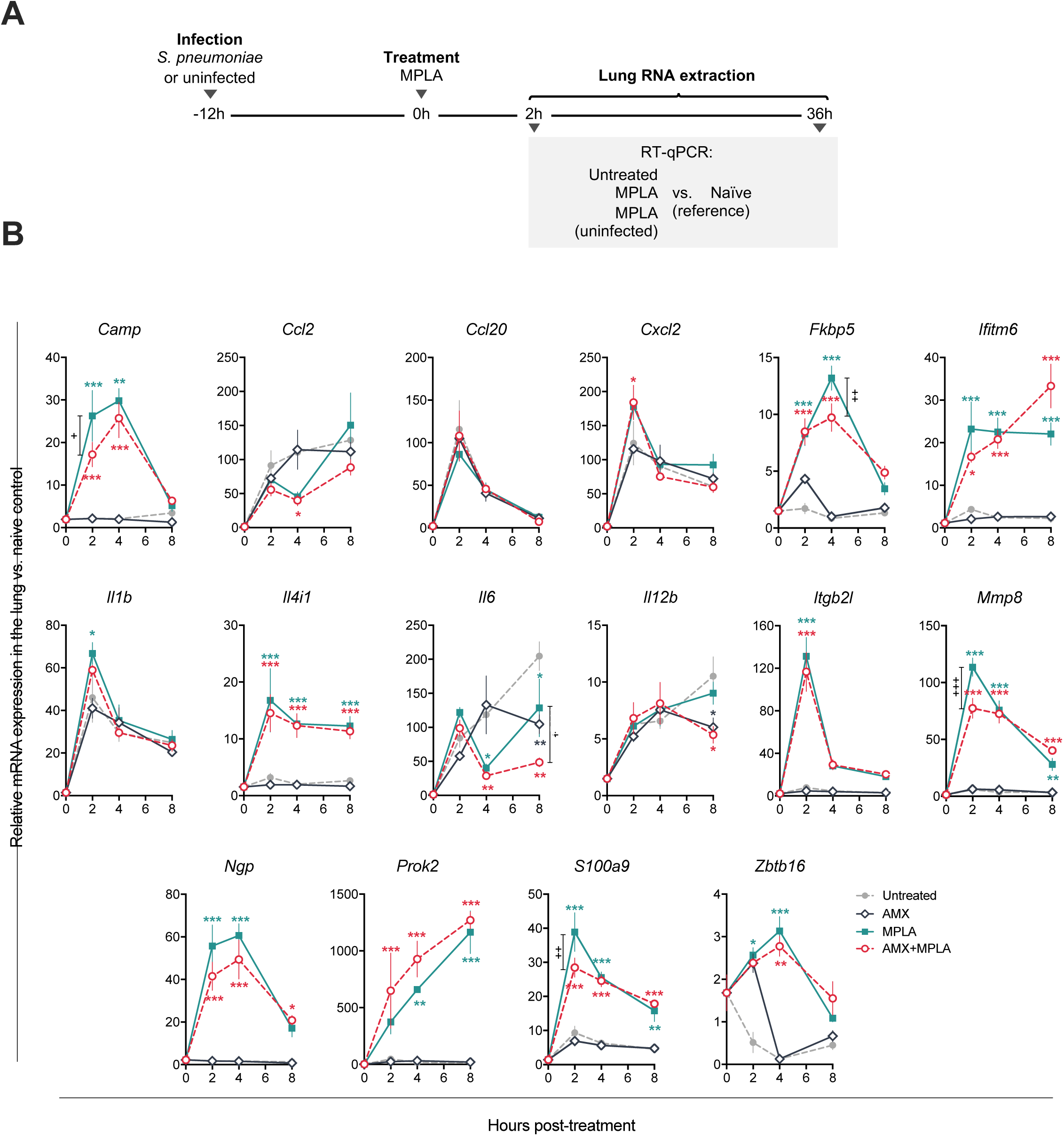
MPLA treatment can accelerate infection-dependent immune responses at the infection site. (A) BALB/c mice (n=3-8/group) were infected with *S. pneumoniae* and treated 12 hours post-infection with either 10 µg of AMX intragastrically, 50 µg MPLA intraperitoneally, a combination of the two treatments (AMX+MPLA), or left untreated; total RNA was extracted from lungs collected at different time points, and mRNA transcripts were compared using RT-qPCR assays. Expression levels were normalized against uninfected, untreated (naïve) controls. (B) Change in relative mRNA expression levels over time for selected genes under different treatment conditions. The mean ± SEM values are shown. A two-way ANOVA with Bonferroni’s post-test for multiple comparisons was applied. *=p<0.05, **=p<0.01 and ***=p<0.001 vs. the mock control group; +=p<0.05, ++=p<0.01, and +++=p<0.001 vs. the indicated comparator groups.

### The systemic response to AMX+MPLA combination treatment during pneumonia

Since combination treatment had significantly outperformed AMX and MPLA monotherapies by limiting bacterial dissemination, we further investigated the effects of treatment on the systemic compartment. In accordance with the gene expression patterns observed in the lungs, we found that the administration of MPLA (whether concomitant with AMX treatment or not) resulted in the sharp release of pro-inflammatory mediators (such as IL-12 p40, IL-6, and CCL2) into the blood (**Figure 5**). The serum concentrations of these cytokines peaked within the first two hours of administration, after which time they fell gradually and returned to baseline levels within six to eight hours. It is noteworthy that during the first 12 hours post-treatment, neither mock-nor AMX-treated infected animals appeared to produce this type of cytokine response - further suggesting that an MPLA-dependent immediate cytokine response is instrumental in better controlling the systemic spreading of bacteria. Although the MPLA- and AMX+MPLA-treated animals displayed similar blood levels of inflammatory mediators, the survival rate was significantly higher in AMX+MPLA group – thus highlighting the importance of the antibiotic’s contribution to the therapeutic efficacy of combination treatment. Since our attempts to measure MPLA in serum were not successful, we assumed that systemic as well as lung immune responses were surrogate markers of MPLA’s effect and thus a way to quantify the pharmacodynamics (PD).

**Figure 5.**
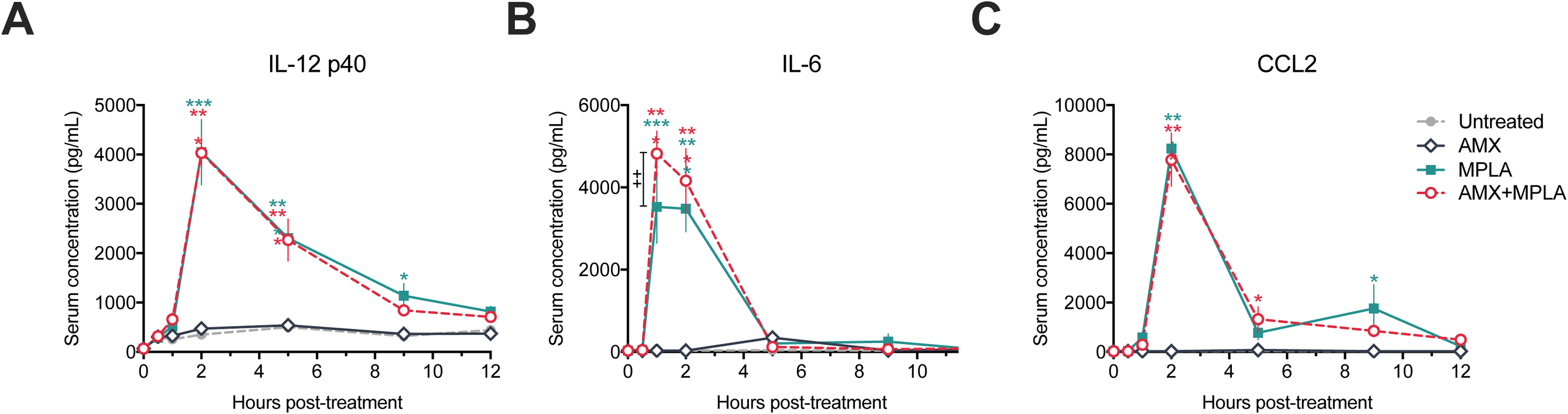
MPLA treatment induces an immediate, transient cytokine response. CD-1 mice (n=4/group) were infected intranasally with 1×10^6^ *S. pneumoniae* and then treated 12 hours later with either AMX (10 µg, intragastric administration), MPLA (50 µg intraperitoneal administration), a combination of AMX and MPLA, or left untreated. Blood samples were collected at different time points post-treatment. Serum levels of pro-inflammatory mediators were determined using ELISAs. (A) IL-12 p40 subunit, (B) IL-6, and (C) CCL2. The mean ± SEM values are shown. A two-way ANOVA with Bonferroni’s post-test for multiple comparisons was applied. At indicated time points: *=p<0.05, **=p<0.01 and ***=p<0.001 vs. the untreated group; ++=p<0.01 vs. the indicated comparator groups.

We observed a typical pharmacokinetic (PK) profile after the oral administration of 10 µg AMX per mouse, with a rapid increase and a rapid subsequent decline in the serum concentration of AMX (**Figure 6**). Interestingly, we observed that the antibiotic’s maximum serum concentration and rate of decline in the AMX+MPLA treatment group were slightly but significantly different from those recorded in the AMX group. Taken as a whole, these findings suggest that MPLA and AMX’s particular effects and different PK profiles in the systemic compartment may contribute to the observed efficacy of the combination treatment.

**Figure 6.**
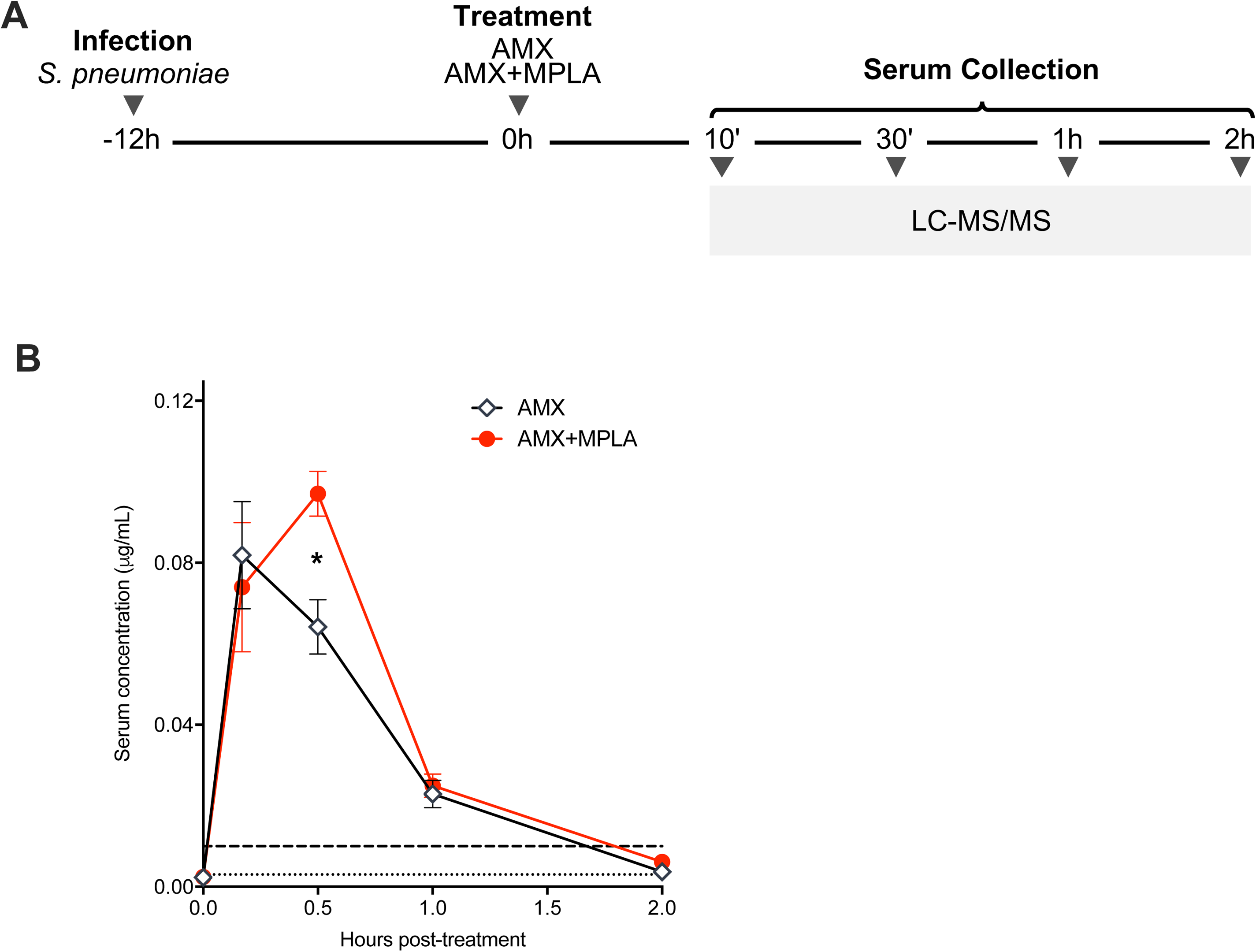
Pharmacokinetics of orally administered AMX in mice. (A) CD-1 mice (n=4/group) infected with 1×10^6^ *S. pneumoniae* and treated with either 10 µg intragastrically administered of amoxicillin (AMX) or a combination of AMX and a 50 µg intraperitoneally administered MPLA (AMX+MPLA). Blood samples were collected at different time points post-treatment. (B) Change in the serum AMX concentration over time, as determined using LC-MS/MS. A two-way ANOVA with Bonferroni’s post-test for multiple comparisons was applied. The values shown are geometric means + range; *=p<0.05 vs. the indicated comparator groups at indicated time points. The dashed and dotted lines along the x-axis indicate the lower limit of quantification (0.01 µg/mL) and the limit of detection (0.003 µg/mL), respectively.

## DISCUSSION

The growing incidence of AMR threatens to limit the currently available treatment options for many bacterial infections. In the present study, we described an alternative strategy for combating invasive pneumococcal disease in an experimental model. We used an immunomodulator (the TLR4 agonist MPLA) as an add-on treatment to boost the efficacy of first-line antibiotic therapy. Our findings confirmed that the outcomes of sub-curative antibiotic treatment are significantly improved (i.e. greater bacterial clearance and better tissue recovery) following targeted stimulation of the host’s innate immune system.

It has already been shown that prophylactic administration of MPLA confers protection against bacterial infection in sepsis, pneumonia, and burn-wound models of disease (15, 17, 18, 22). In our mouse model of progressive pneumococcal pneumonia, we observed that the single-shot, systemic, post-infection administration of MPLA was associated with a significantly lower bacterial load in lung and spleen and higher survival rates. These observations demonstrated that MPLA could potentially function as a therapeutic agent. On the same lines, a recent study evidenced the therapeutic effect of MPLA-containing adjuvants in the context of systemic mycosis (19). Another study reported that administration of LPS, the highly pyrogenic and toxic TLR4 agonist with antibiotic is able to eliminate Salmonella invading the mesenteric lymph nodes, in contrast to stand-alone antibiotic treatment (23). Together with our present findings, these literature data suggest that MPLA could be repurposed as a universal antimicrobial drug whose therapeutic activity is independent of the infection route and the microbial pathogen. Despite MPLA’s proven adjuvant potency, a single dose of MPLA alone had a limited therapeutic effect. We therefore hypothesized that MPLA’s immunostimulatory properties could be best exploited as a non-specific and safe booster of innate immune responses (thus improving the efficacy of an otherwise suboptimal dose of antibiotic), rather than as a direct antimicrobial treatment.

In the present proof-of-concept study with an AMX-susceptible strain of *S. pneumoniae*, we sought to replicate clinical conditions. Firstly, we treated the mice with a low dose of AMX; this mimics the context of AMR in which a poorly administered or incorrectly dosed antibiotic treatment leads to incomplete bacterial clearance or the development of resistance. Secondly, we gave the bacteria time to establish an infection prior to treatment; this simulated both the lag between diagnosis and treatment, and the cascade of immune responses to the initial bacterial insult. The combination of AMX and MPLA in a single administration had a greater therapeutic effect than each monotherapy alone. It is known that *S. pneumoniae* possesses a large number of virulence factors that facilitate colonization. These include cell wall peptidoglycan components and pneumolysin, both of which appear to elicit MyD88-dependent immune responses via TLR2- and TLR4-specific pathways, respectively (24-27). Furthermore, TLR9 has been shown to have a major role in early host defenses against *S. pneumoniae* infections (28, 29). Given that MPLA immunostimulation is biased towards TRIF-mediated signaling downstream of TLR4, it is tempting to speculate that the introduction of *S. pneumoniae* provides just the right type and right amount of primary stimulation via MyD88, which is then amplified by the introduction of the secondary stimulus (MPLA) - leading to an enhanced immune response through the activation of both MyD88- and TRIF-dependent pathways. Toll-like receptor-specific priming induced by pneumococcal pneumonia may be a major factor in the enhanced immune response observed upon MPLA treatment in our model.

It remains to be seen how MPLA is involved in greater bacterial clearance in the context of treatment with sub-curative doses of AMX. The MPLA-associated transcriptional signature in the lungs of infected animals highlighted the upregulation of genes associated with neutrophil function and tissue homing. One can assume that a greater neutrophil count and an enhanced killing capacity may drive MPLA’s antibacterial action and synergize them with the antibiotic’s effects in the airways. Our preliminary data at 12h post-treatment did not highlighted changes in neutrophil number in lung and spleen; however, monocytes were found in higher number in lung (**Figure S4**). Similarly, the systemic effects of MPLA-mediated signaling (such as the transient production of cytokines and chemokines) may link immune cell activation to bacterial clearance in the peripheral tissues. Our previous research on the TLR5 agonist flagellin identified two contributory mechanisms in bacterial clearance: (i) greater infiltration of myeloid cells into the lungs and airways, and (ii) the activation of IL-17/IL-22 responses by innate lymphoid cells (20, 21, 30, 31). Here, the lung transcriptional signature suggests that MPLA influences the myeloid cell compartment by promoting recruitment and activating antibacterial activity. In contrast to flagellin, MPLA administration was not associated with the rapid production of IL-17/IL-22 by lung innate lymphoid cells; this is also the case for LPS (31). TLR signaling is associated with broad range of antibacterial mechanisms in various cell types, some of which are probably independent of AMR mechanisms (4). A multifactorial action is also a means of developing a therapeutic approach that is less likely to promote AMR. With a view to defining specific immune targets, it will be important to determine how the MPLA-stimulated immune system cooperates with an antibiotic to improve treatment efficacy.

Drug PK and PD studies provide a quantitative basis for dosing regimens in humans and other animals. Orally administered AMX is absorbed rapidly in humans, and has a relatively short period of serum availability (32-34). The absorption in mice is reportedly even faster (35, 36). The PK characteristics of AMX indicate that this antibiotic is efficacious very soon after administration. Hence, AMX’s bactericidal activity depends greatly on the time during which the serum concentration exceeds the minimum inhibitory concentration for its target; accordingly, multiple, regular administrations are required to ensure efficacy (37, 38). Although these treatment regimens are used widely, they are always associated with the risk of encouraging the development of AMR. Amoxicillin’s rapid PK maximum and minimum concentrations were in contrast with the PD characteristics of MPLA, i.e. longer-term activation of the immune system. Our present findings suggest that the different time scales of AMX’s and MPLA’s respective activities increased the overall efficacy of treatment when the two are administered together. On a PK level, we also observed a slight increase in AMX retention when latter was combined with MPLA. Future research must focus on whether and how the two substances interact specifically *in vivo* in an infectious context.

Our present results highlighted on the potential of targeting innate immunity with a TLR4 agonist as a viable strategy for improving antibiotic therapy against a bacterial infection. This pragmatic approach uses the host’s immune system to strengthen the attack against invading microorganisms, and may also help to repurpose currently available drugs with known characteristics. Our experimental evidence suggests that the enhanced therapeutic effect of the MPLA-AMX combination is achieved through a combination of TLR4 priming and the time scales of MPLA’s and AMX’s respective peak biological activities. In the future, it will be important to investigating possible synergistic relationships between individual treatment components by using (i) PK/PD analyses and comprehensive mathematical modelling, and (ii) further *in vivo* simulations in multidrug resistant strains treated with high-dose and/or multiple-dose antibiotic regimens. Ultimately, this promising approach may open up new avenues for the design of host-directed therapeutics for infectious diseases.

## MATERIALS AND METHODS

### Ethics statement

Animals were maintained in individually ventilated cages and were handled in a vertical laminar flow cabinet (biosafety level 2). All experiments complied with current national, institutional and European regulations and ethical guidelines, were approved by our Institutional Animal Care and Use Committee (animal facility agreement C59-350009, Institut Pasteur de Lille; reference: APAFIS#5164, protocol 2015121722429127_v4) and were conducted by qualified, accredited personnel.

### Bacterial strains and cell cultures

*Streptococcus pneumoniae* serotype 1 (clinical isolate E1586) was obtained from the Uruguayan Ministry of Health’s National Reference Laboratory (Montevideo, Uruguay). Working stocks were prepared as described previously (20, 21). Briefly, Todd Hewitt Yeast Broth (THYB) (Sigma-Aldrich, St. Louis, MO, USA) was inoculated with fresh colonies grown in trypcase soy agar plates supplemented with 5% sheep blood (BioMérieux, Marcy-l’Étoile, France), and incubated at 37°C until the OD_600nm_ reached 0.7-0.9 units. Cultures were stored at -80°C in THYB + glycerol 12% (vol/vol) for up to 3 months. For mouse infections, working stocks were thawed and washed with sterile Dulbecco’s phosphate-buffered saline (PBS; Gibco, Grand Island, NY, USA) and diluted to the appropriate concentration. The number of bacteria (expressed in colony forming units [nezs]) was confirmed by plating serial dilutions onto blood agar plates.

### The mouse model of infection

Six-to eight-week-old female BALB/cJRj (BALB/c), C57BL/6JRj (C57BL/6) or RjOrl:SWISS (CD-1) mice (Janvier Laboratories, Saint Berthevin, France) were used for all *in vivo* experiments. Infection was carried out as described previously (30, 39). Briefly, mice were first anesthetized by intraperitoneal injection with a solution of 1.25 mg ketamine plus 0.25 mg xylazine in 250 µL of PBS, after which they were infected intranasally with a 30 µL PBS suspension containing 1 to 4 × 10^6^ CFU of *S. pneumoniae*. All the treatments described below were administered once 12h post-infection. Mice were sacrificed at selected times via the intraperitoneal injection of 5.47 mg of sodium pentobarbital in 100 µl PBS (Euthasol, Virbac, Carros, France). Blood was sampled by retro-orbital puncture into Z-Gel micro tubes to prepare serum for downstream applications (Sarstedt, Nümbrecht, Germany). Lungs and spleens were collected, homogenized, and plated onto blood agar to determine the endpoint bacterial load. For survival assays, both mortality and changes in body weight were monitored; mice were individually weighed prior to infection and then every 24h for a period of up to two weeks.

### Reagents and administration of treatments *in vivo*

Amoxicillin trihydrate (Sigma-Aldrich) was prepared in a stock solution of 1.75 mg/mL in sterile water and then adjusted to a final dose of 5, 30, or 350 µg/mouse (i.e., 0.2, 1.2, or 14 mg/kg) before intragastric administration of a volume of 200 µL by oral gavage. Lipopolysaccharide from *E. coli* O111:B4 (S-form) and MPLA from *S. minnesota* R595 (Re) (TLR*pure*™, Innaxon Therapeutics, Bristol, United Kingdom) were obtained as sterile solutions or prepared from powder with sterile distilled water to a concentration of 1 mg/L (according to the manufacturer’s recommendations) and then adjusted to different final concentrations in sterile PBS and administered by intraperitoneal injection (200 µL).

### Quantification of gene expression and microarrays

Lungs were perfused with PBS prior to sampling. Lung or liver total RNA was extracted with the Nucleospin RNA II kit (Macherey Nagel, Düren, Germany) and reverse-transcribed with the High-Capacity cDNA Archive Kit (Applied Biosystems, Foster City, CA, USA). The cDNA was amplified using SYBR-Green-based real-time PCR on a Quantstudio™ 12K Real-Time PCR System Thermo Fisher Scientific, Carlsbad, CA, USA). Specific primers used are listed in **Table S1**. Relative mRNA levels were determined by comparing the PCR cycle thresholds (Ct) for the gene of interest vs. *Actb* (ΔCt) and then the ΔCt values for treated vs. untreated (mock) groups (ΔΔCt). For the microarray analysis, total RNA yield and quality were further assessed on the Agilent 2100 bioanalyzer (Agilent Technologies, Santa Clara, CA, USA). One-color whole mouse (084809_D_F_20150624 slides) 60-mer oligonucleotide 8×60k v2 microarrays (Agilent Technologies) were used to analyze gene expression. The cRNA labelling, hybridization and detection steps were carried out according to the supplier’s instructions (Agilent Technologies). For each microarray, cyanine-3-labeled cRNA was synthesized from 50 ng of total RNA using the low-input QuickAmp labeling kit. RNA Spike-In was added to all tubes and used as a positive control in the labelling and amplification steps. Next, 600 ng of each purified labelled cRNA were then hybridized and washed following manufacturer’s instructions. Microarrays were scanned on an Agilent G2505C scanner, and the data were extracted using Agilent Feature Extraction Software (version 10.7.3.1, Agilent Technologies). Microarray data have been deposited in the Gene Expression Omnibus database (accession number: GSE118860). Statistical comparisons and filtering were performed with the Limma R package with 75-percentile normalization. Differentially expressed genes were considered to be those with an adjusted p-value below 0.05 after the false discovery rate had been checked with the Benjamini-Hochberg procedure (40). Pathways were investigated using Ingenuity Pathway Analysis software (Ingenuity Systems, Redwood City, CA, USA).

### Histology

Lungs were fixed by intratracheal perfusion with 4% formaldehyde prior to sampling. The left lobe and the upper right lobe were included in paraffin, and 3-to 5-µm tissue sections were stained with hematoxylin and eosin reagent. The slides were blindly evaluated for neutrophil infiltration, perivascular infiltration, edema, and pleuritis on a 6-level scale, where 0 corresponded to the absence of lesions, and 1 to 5 corresponded to minimal, slight, moderate, marked, and severe lesions, respectively (Althisia, Troyes, France).

### Quantification of serum cytokine levels

Serum levels of CCL2, IL-6 and IL-12 p40 were measured using an ELISA, according to the manufacturer’s instructions (R&D Systems, Minneapolis, MO, USA).

### Determination of serum amoxicillin concentrations

Serum AMX concentrations were assayed using previously developed and validated liquid chromatography tandem mass spectrometry (LC-MS/MS) method (41). In brief, the proteins in 10 µL of serum were precipitated with 40 µL of ice-cold methanol. After diluting the supernatant with water, the sample was injected into the LC system (Agilent Technologies) by using a gradient elution at a flow rate of 0.3 mL/min with acetonitrile and water with formic acid. The AMX ion product (*m/z* 114) was quantified using electrospray ionization MS in positive ion mode over a calibration range from 0.01 to 10 µg/mL. In-study validation was performed according to the European Medicines Agency guidelines on bioanalytical method development (42)

### Statistical analysis

Results were expressed as the median and individual values, median [interquartile range] or mean ± standard error of the mean (SEM), as appropriate. Groups were compared using a Mann-Whitney test (for two independent groups) or a Kruskal-Wallis one-way analysis of variance (ANOVA) with Dunn’s post-test (for three or more groups). The log rank test was used for survival analyses. Statistical analyses were performed using GraphPad Prism software (version 8.2, GraphPad Software Inc., San Diego, CA, USA), and the threshold for statistical significance was set to p<0.05.

## ACKNOWLEDGMENTS

We thank Shéhérazade Sebda and Frédéric Leprêtre from University of Lille’s genomics facility for preparing microarrays and depositing the data in the Gene Expression Omnibus database, respectively.

## AUTHOR CONTRIBUTIONS

FC performed all animal, RT-qPCR, ELISA, and flow cytometry experiments. SF and RM analyzed antibiotic PK data. LM provided FC with technical assistance. MF performed microarray experiments and bioinformatics analyses. CK, CC, and JCS designed the experiments. FC, JCS, and CC wrote the manuscript. JCS and CC supervised the experimental work as a whole.

## DATA AND MATERIALS AVAILABILITY

Microarray data are available in the Gene Expression Omnibus database (accession number: GSE118860). The raw data supporting the conclusions of this manuscript will be made available by the authors, without undue reservation, to any qualified researcher.

